# The dynamics of preferential host switching: host phylogeny as a key predictor of parasite prevalence and distribution

**DOI:** 10.1101/209254

**Authors:** Jan Engelstädter, Nicole Z. Fortuna

## Abstract

This preprint has been reviewed and recommended by *Peer Community In Evolutionary Biology* (https://doi.org/10.24072/pci.evolbiol.100049).

New parasites commonly arise through host-shifts, where parasites from one host species jump to and become established in a new host species. There is much evidence that the probability of host-shifts decreases with increasing phylogenetic distance between donor and recipient hosts, but the consequences of such preferential host switching remain little explored. We develop a mathematical model to investigate the dynamics of parasite host-shifts in the presence of this phylogenetic distance effect. Host trees evolve under a stochastic birth-death process and parasites co-evolve concurrently on those trees, undergoing host-shifts, co-speciation and extinction. Our model indicates that host trees have a major influence on these dynamics. This applies both to individual trees that evolved under the same stochastic process and to sets of trees that evolved with different macroevolutionary parameters. We predict that trees consisting of a few large clades of host species and those with fast species turnover should harbour more parasites than trees with many small clades and those that diversify more slowly. Within trees, large clades should exhibit a higher infection frequency than small clades. We discuss our results in the light of recent cophylogenetic studies in a wide range of host-parasite systems, including the intracellular bacterium *Wolbachia*.

## Introduction

Parasitism represents one of the most successful modes of life. Humans harbour more than 1400 species of parasites (Taylor *et al.* 2001), which extrapolates to an enormous total number of parasites across all host species. Where do all these parasites come from? Some parasites may already have been present in their host species’ ancestor and maintained ever since. This scenario of ‘cospeciation’ has been described in some mutualists but appears to be rare in parasites (de Vienne *et al.* 2013). Other parasites may originate from organisms that are either free-living, or non-parasitic symbionts (Crook 2014; Hurst 2016). Finally, some parasites may have switched from another host species to their present-day host. Such host-shifts have been widely documented. The majority of human pathogens originate through host-shifts, including HIV and malaria (Wolfe *et al.* 2007). Host-shifts are also the predominant cause of new host-parasite associations for *Wolbachia* endosymbionts and their arthropod hosts (Werren *et al.* 1995), rabies viruses in bats (Streicker et al. 2010), lentiviruses in primates (Sharp *et al.* 2000), oomycetes in Asteraceae (Choi & Thines 2015), and malaria in birds (Ricklefs *et al.* 2014).

Establishing a sustainable relationship with a new host species represents a considerable challenge to parasites. While many opportunities for host-switches exist, most attempts are unsuccessful and lead to mere ‘spill-over’ infections, i.e. infections with no or short transmission chains (Taylor *et al.* 2001; Wood *et al.* 2012). Examples of such spillovers in humans include rabies, Hendra, and Ebola viruses. Successful host-shifts are difficult because the parasite must be able to enter, proliferate within, and transmit efficiently between, members of a new host species that they are not adapted to. These requirements mean that all else being equal, shifts to new hosts that are similar to the original host with respect to relevant traits should be easier than shifts to hosts that are very different from the original one. Given that this similarity will be positively correlated with phylogenetic relatedness between host species, we can predict that host-shifts to closely related new hosts should be more common than host-shifts to distantly related hosts (Charleston & Robertson 2002; Engelstädter & Hurst 2006; Longdon *et al.* 2014). We will refer to this expectation as the ‘phylogenetic distance effect’.

There are two lines of evidence for the phylogenetic distance effect. First, a number of transfection experiments have been conducted in which parasites from one species were exposed to a range of hosts from different species. For example, Longdon *et al*. (2011) demonstrated that for three sigma viruses endogenous to different species of *Drosophila*, phylogenetic distance between the donor and recipient host species was negatively correlated with the viruses’ ability to replicate within the recipient host. Similarly, for male-killing *Spiroplasma* bacteria in ladybird beetles, Tinsley & Majerus (2007) reported that as the distance between the original host and a new host increased, the ability of the parasite to kill male offspring (the phenotype driving the infection) was reduced. Other systems in which experimental evidence for the phylogenetic distance effect has been obtained include nematodes infecting *Drosophila* flies (Perlman & Jaenike 2003), feather-lice infecting pigeons and doves (Clayton *et al.* 2003), and plant-fungal systems (Gilbert & Webb 2007; de Vienne *et al.* 2009). Strong evidence for the phylogenetic distance effect from 25 publications reporting the success or failure of *Wolbachia* transfection experiments is reviewed in Russell *et al*. (2009).

Second, different phylogenetic methods have been used to investigate whether host-shifts occur preferentially between related host species. Much early work comparing host and parasite phylogenetic trees focused on reconciling those trees and identifying the degree of cospeciation. However, Charleston & Robertson (2002) showed that the observation that closely related lentiviruses tend to infect closely related primate hosts is best explained not by codivergence but by preferential host-switching between related hosts (because the viruses only spread relatively recently on the primate tree). Studies of rabies viruses infecting various bat species confirmed the presence of the phylogenetic distance effect (Streicker *et al.* 2010) and further demonstrated that while species range overlap was the best predictor of spillover events, phylogenetic distance was the best predictor of host-shift events (Faria *et al.* 2013). Clark & Clegg (2017), studying the distribution of malaria among south-Melanesian birds, found that despite ample opportunity for host-switching due to vector-borne transmission, similar parasites were restricted to similar hosts. Some studies have also provided evidence that *Wolbachia* endosymbionts switch preferentially between related arthropod host species (Baldo *et al.* 2008; Russell *et al.* 2009; see also Discussion). In summary, the experimental and comparative work indicates that although not ubiquitous (e.g., Stahlhut *et al.* 2010; Longdon *et al.* 2015), the phylogenetic distance effect is an important determinant of host-shifts in many systems.

Most of the previous theoretical work on host-shifts has focused on reconciling host and parasite phylogenetic trees, identifying host-shift vs. cospeciation events, and inferring parameters underlying these processes (older literature reviewed in de Vienne *et al.* 2013; newer work includes Baudet *et al.* 2015; Wieseke *et al.* 2015; Drinkwater & Charleston 2016; Alcala *et al.* 2017). Mathematically speaking, these are very hard problems and most of the developed algorithms are computationally expensive. It is therefore not surprising that the phylogenetic distance effect is usually not considered in these methods, despite the widely recognised fact that preferential host switching may be misinterpreted as cospeciation (de Vienne *et al.* 2007). Exceptions include a study where data from RNA virus-mammal associations were used to test two different models describing the decline in host-shift success with increasing phylogenetic distance between host species (Cuthill & Charleston 2013), and a study in which the host-shift dynamics of protozoan parasites in new world monkeys were inferred (Waxman *et al.* 2014). In contrast to the development of inference methods for host-parasite cospeciation and host-shifts, little work has been done to explore the consequences of the phylogenetic distance effect for the dynamics of parasites spread between host species and the expected patterns of parasite distributions. In simulations of parasite host switching, Engelstädter and Hurst (2006) demonstrated that the ‘shape’ of a host clade strongly influences parasite prevalence and distributions within host clades. However, their model (like the model by de Vienne *et al.* 2007) only considered a few idealised host trees (e.g., either completely symmetrical or ladder-like), and they (like Cuthill & Charleston 2013; Waxman *et al.* 2014) assumed that host switching occurred only at the tips of the trees.

Here, we present the results of a stochastic model in which a clade of host species evolves under a birth-death process and a clade of parasites spreads concurrently on this host tree through both cospeciation events and host-shifts (either preferential or random). Through extensive computer simulations we investigate how often the parasites can invade a naïve host tree, how many hosts will become infected and how the parasites are distributed across host species. Our model predicts that both individual host phylogenies and the macroevolutionary processes underlying these phylogenies have a major influence on host-shift dynamics when the phylogenetic distance effect is important.

## Methods

### Mathematical model

We considered a stochastic model of host-parasite co-diversification, illustrated in Figure 1. Host trees emerge from a single ancestor according to a density-dependent birth-death process. Hosts go extinct at a constant rate μ and speciate at a baseline rate *λ* that is multiplied by the term (1-*N*/*K*), resulting in a decreasing speciation rate as the number of host species *N* approaches the carrying capacity *K*.

**Figure 1.**
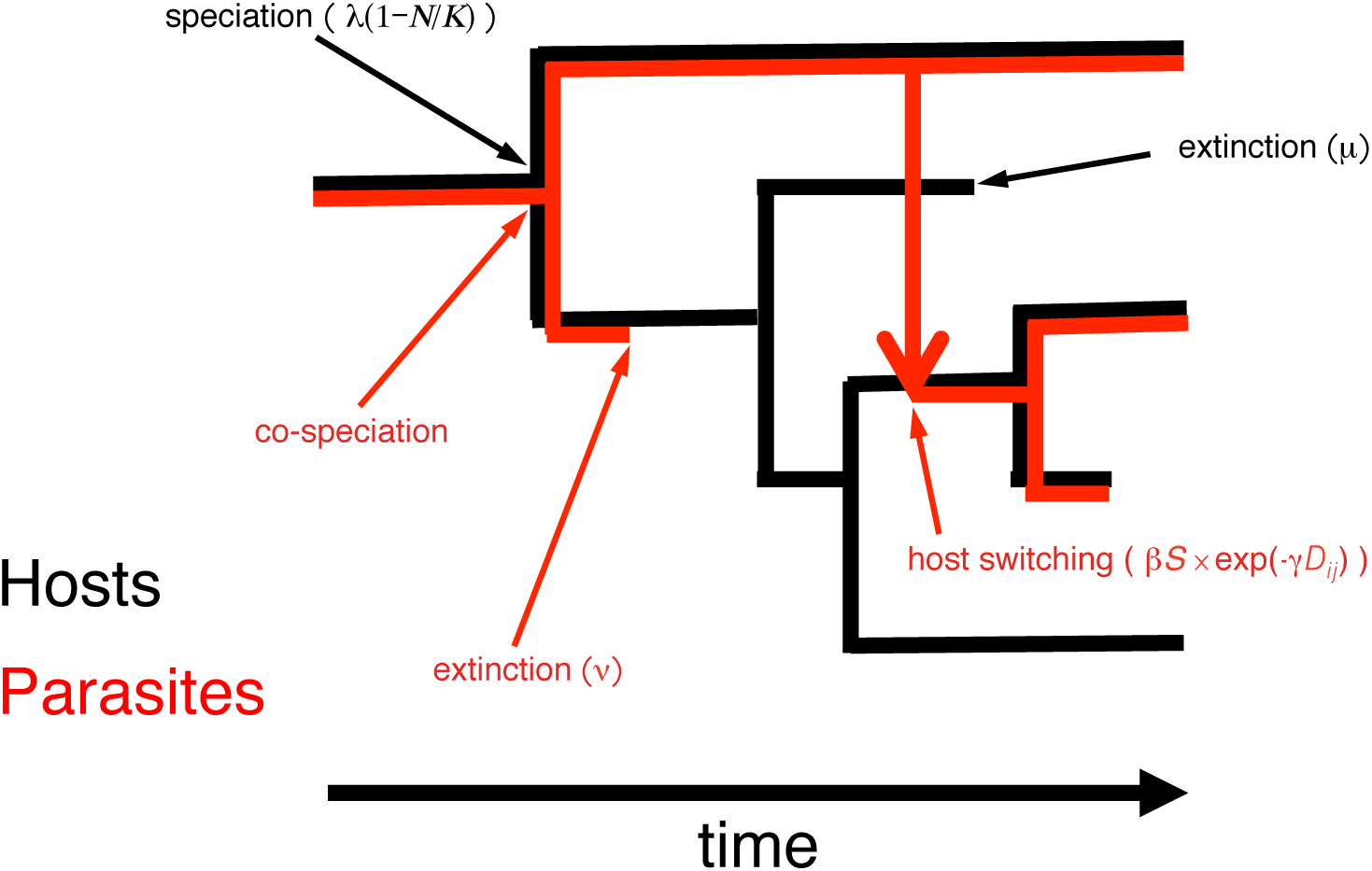
Illustration of the model; see Methods section for details.

Each parasite species is associated with a single host species. Parasites go extinct at a constant rate *v* and always co-speciate whenever their hosts speciate. Host-shifts represent an alternative, independent mode of parasite speciation in which one lineage remains associated with the original host and a new lineage arises that is associated with a new host species. Host-shifts occur at a baseline rate β(*N*-1) per parasite lineage. Potential new hosts are chosen randomly but not all host-shifts are successful. First, host-shifts are unsuccessful if the new host is already infected (but see below for an extension of the model where this assumption is relaxed). Second, parasites may not become established if the new host is phylogenetically too distant from the original one. Specifically, we assume a parasite establishment probability, exp(-*γD_ij_*). (the same relationship but using a different notation was used by Engelstädter & Hurst 2006; Cuthill & Charleston 2013). Here, the parameter *γ* determines how fast the establishment probability declines with increasing phylogenetic distance *D*_*ij*_ between the donor host species *i* and the new host species *j* (i.e., *D*_*ij*_ is the total length of branches connecting the two species with their most recent common ancestor). When *γ*=0, all host-shifts are successful (no phylogenetic distance effect) but with larger values of *γ*, host species that are distantly related to the original host are increasingly unlikely to become infected.

In addition to this basic model, we also investigated three model extensions that incorporate 1) coinfection of multiple parasites in one host species, 2) parasite loss during cospeciation, and 3) within-host speciation of parasites. For details we refer to the Supplementary Information (SI), section 1.

### Model implementation

We analysed our model using computer simulations. Time proceeds in small steps (Δt=10^−4^) in which the different events (host speciation, host extinction etc.) take place with probabilities given by their rates multiplied by Δt. Since host evolution is not affected by the parasites in our model, we first simulated the host trees and then simulated parasite diversification on those host trees.

The routines to simulate the cophylogenetic process were implemented in the programming language R (R Core Team 2017). We bundled these routines, along with other functions for simulation, subsequent analyses and plotting of cophylogenetic trees, into a new R-package named ‘cophy’. This package depends on the R-packages ape v4.1 (Paradis *et al.* 2004), parallel v3.3.2 (R Core Team 2017), foreach v1.4.3 (Revolution Analytics & Weston 2015b), and doParallel v1.0.10 (Revolution Analytics & Weston 2015a). We used the R-packages devtools v1.13.2 (Wickham & Chang 2017) and roxygen2 v6.0.1 (Wickham *et al.* 2017) to generate our package. For data analysis, we also used lme4 v1.1-12 (Bates *et al.* 2015) and vegan v2.4-5 (Oksanen et al. 2017). The cophy package is available on GitHub at https://github.com/JanEngelstaedter/cophy/tree/master/cophy.

### Simulations

We started by simulating different sets of host trees, each containing 100 trees that were initialised with a single species and evolved for 100 time units. Only trees that survived this time span were retained. For an initial standard set of trees, we chose a speciation rate of *λ*=1, an extinction rate of μ=0.5 and a carrying capacity of *K*=200, yielding an expected equilibrium tree size of *N*=100 species. Using this set as a baseline, we created three series of similar sets with 1) the same speciation and extinction rate but with *N* increasing from 30 to 200, 2) the same equilibrium clade size and net diversification rate (*λ*–μ=0.5), but extinction rate μ increasing from 0.1 to 0.9, and 3) eight other sets with the same equilibrium clade size but different net diversification and turnover rates (see SI section 2.1 for details).

To simulate parasite diversification on those host trees, we introduced a single parasite species at time *t*=50 on a given host tree and simulated until the parasite went extinct or the present (*t*=100) was reached. For each host tree, we randomly chose ten branches on which the first parasite species arrived and performed ten replicate simulations for each of these initial branches. Thus, for each set of host trees we performed a total of 100×10×10=10,000 simulations.

We focused on two parameter sets for parasite evolution. First, we used a parameter combination with which the phylogenetic distance effect is present: β=0.5, *γ*=0.06 and *v*=1. Second, as a control, we used a parameter combination with which the phylogenetic distance effect is absent: β=0.02, *γ*=0 and *v*=1. We refer to these two standard parameter combinations as the standard PDE and no-PDE parameters, respectively. The parameters were chosen so that both the probability of parasite establishment and the observed frequency of infected hosts at the end of the simulation are roughly the same (around 0.5; see Results). In order to test whether our results are robust with respect to the choice of parameters, we also performed simulations with two other PDE / no-PDE parameter combinations that are characterised by either a lower or a higher turnover rate in parasite diversification. Finally, we also performed the same simulations for our three model extensions (SI section 1).

### Analyses of results

For each simulation we obtained some basic statistics, including the fraction of simulations in which the parasites established a surviving infection on the host trees, the distribution of the number of host and parasite species and the frequency of infected hosts at the end of the simulation (contingent on parasite survival). For parasite trees that did not leave any surviving species we obtained the time of extinction, and for those which did we obtained the time of the most recent common ancestor of all extant species. As a simple statistic describing the distribution of parasites within the host phylogeny we used the correlation coefficient between host and parasite phylogenetic distances (see SI, section 2.2). We also investigated the frequency of infected host species within different clades of the host tree (see SI section 2.3).

## Results

### Patterns of parasite spread and distributions

We first focused on understanding the host-shift dynamics under the phylogenetic distance effect on a standard set of host trees simulated under the same birth-death process. Figure 2 compares some basic summary statistics for simulations in presence vs. absence of the phylogenetic distance effect (standard PDE vs. no-PDE parameters). By choice of parameters, the final mean frequency of infected hosts for simulations with surviving parasites was similar in both cases (Figure 2A). However, the variance in infection frequencies was greater with the phylogenetic distance effect than without (see also below). If the parasites went extinct, this usually occurred early during the simulations in both scenarios (Figure 2B). The most recent common ancestor of all surviving parasites lived later on average with than without the phylogenetic distance effect (Figure 2C), reflecting higher parasite turnover in the latter case.

**Figure 2.**
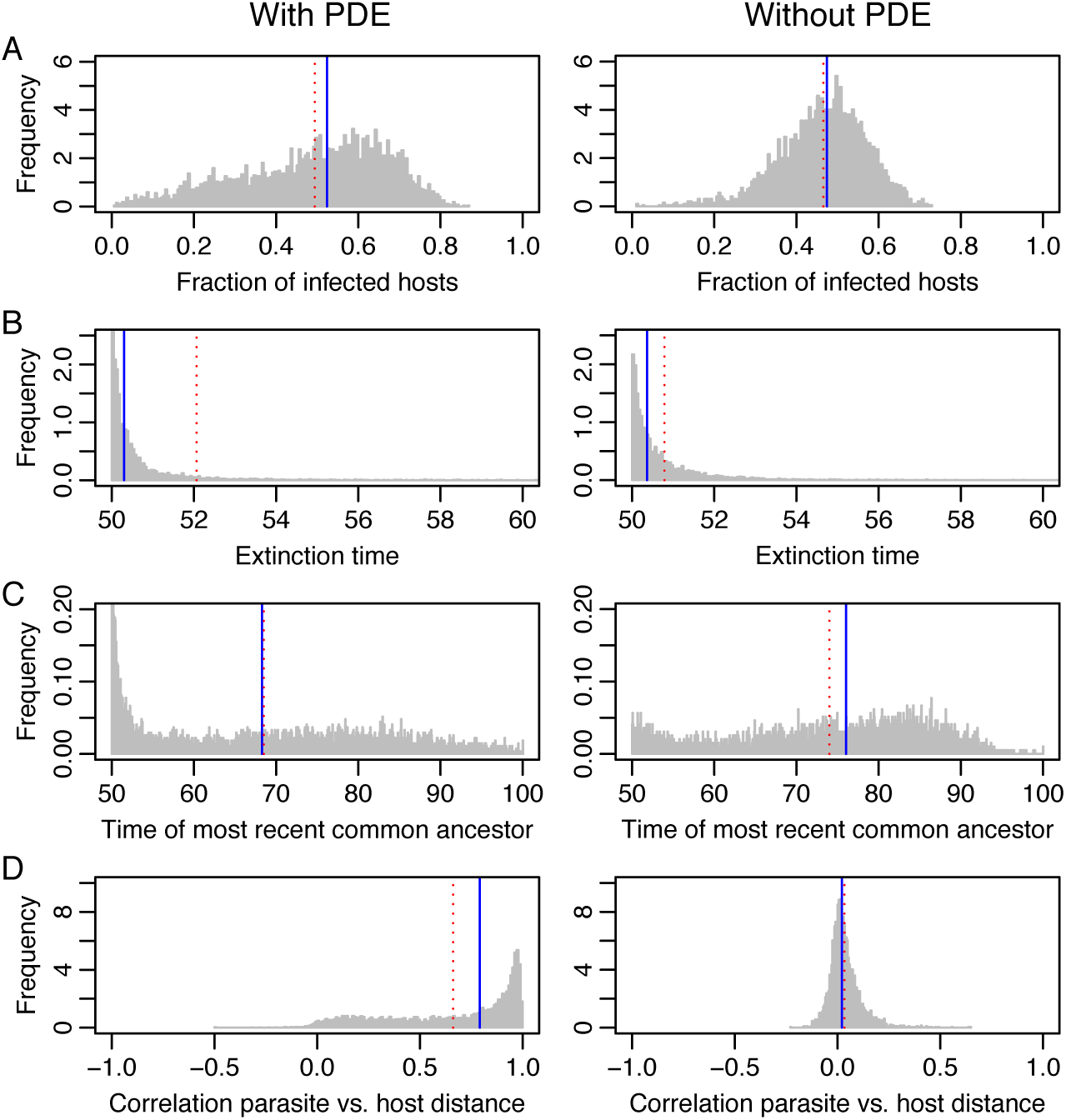
Summary statistics for simulations in the presence and absence of the phylogenetic distance effect, with the standard host tree set and standard PDE vs. no-PDE parameters. Panel (A) shows the distribution of the fraction of infected host species across the 10,000 simulations, contingent on parasite survival. Panel (B) shows the distribution of parasite extinction times when the parasite did not survive following its introduction at time 50. Panel (C) shows the distribution of the time of the most recent common ancestor of all surviving parasite species (where time=100 is the present). In panel (D), the distribution of the correlation between parasite and host phylogenetic distances is shown. In all plots, the solid blue line indicates the median and the dashed red line the mean of the distributions.

In Figure 2D, we plot the distribution of correlation coefficients between phylogenetic distances between pairs of parasite species and the phylogenetic distances between their associated host species. In the presence of the phylogenetic distance effect, this distribution shows a strong positive trend: >98% of simulations where the parasites survived exhibited a positive correlation, with a median of 0.807. Thus, closely related parasites tend to be found in closely related host species and *vice versa*. This is not primarily a consequence of co-speciation events but of the phylogenetic distance effect. In the absence of the phylogenetic distance effect, the host-parasite phylogenetic correlation coefficients are distributed around zero. The median of this distribution is still positive (0.021), which is explained by recent co-speciation events, but the distribution is very distinct from the one observed in the presence of the phylogenetic distance effect.

We can also ask how parasites are distributed within different host clades when the phylogenetic distance effect is important. Parasites will shift predominantly within host clades but rarely between different clades in this case. One might therefore expect that all else being equal, larger host clades should on average harbour more parasites than smaller clades. Figure S1 shows that this expectation is fulfilled both when host trees are split into a few large and into many small clades (Figure S1A and B). In the absence of the phylogenetic distance effect, host clade size has no effect on the fraction of hosts that are infected within those clades (Figures S1C and D).

### Host trees are important in determining parasite spread

Figure 3A shows that in the presence of the phylogenetic distance effect, the distribution of the fraction of infected host species observed at the end of the simulations differs according to host tree. A random effects model confirms the visual impression that much of the variation in the fraction of infected host species observed at the end of the simulations is due to the specific host tree on which the parasites spread (see SI, section 2). By contrast, in the absence of the phylogenetic distance effect, the observed mean infection frequencies are much more homogeneous across host trees (Figure 3B), with a lower fraction of variance explained by host trees (SI section 2).

**Figure 3.**
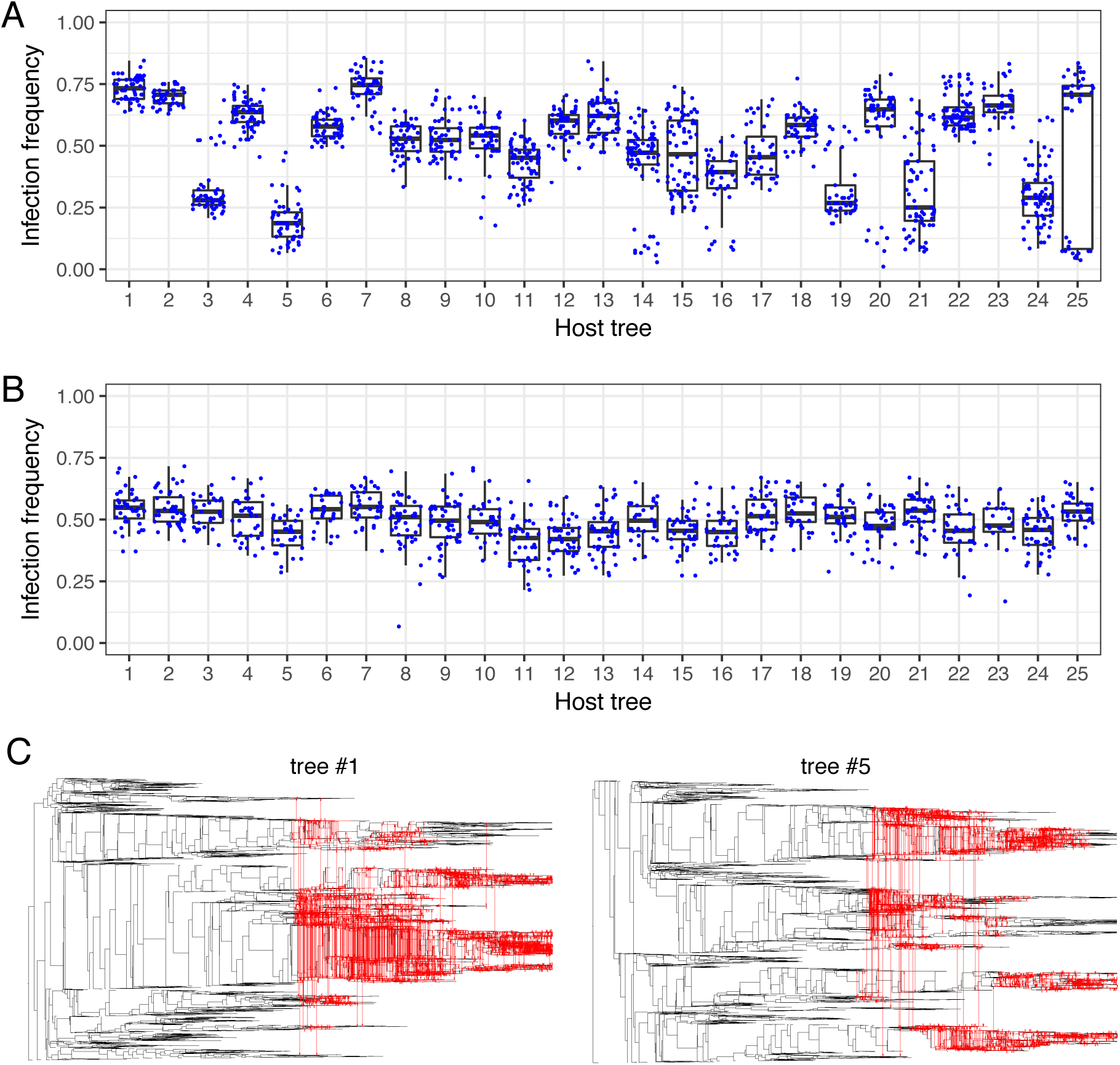
Distributions of infection frequencies with (A) and without (B) the phylogenetic distance effect on the first 25 host trees. Each dot shows the fraction of infected host species at the end of a simulation run. Simulations in which the parasites did not survive until the end of the simulation are not shown. Boxes show the interquartile range with the horizontal line indicating the median and whiskers indicating the distance from the box to the largest value no further than 1.5 times the interquartile range. All parameters take the standard values. Panel (C) shows examples host-shift dynamics for two of the host trees in presence of the phylogenetic distance effect, yielding final infection frequencies of 74% and 24%, respectively. For larger trees and more examples, see Figure S2.

To obtain some intuition for the importance of host trees in shaping host-shift dynamics, consider the example co-phylogenies shown in Figures 3C and S2, corresponding to host trees number 1, 5 and 25. With host tree #1 (Figures 3C and S2A), most of the extant host species form one large, relatively recently formed clade of species. A second, smaller clade is still closely related to the first one. This means that for most host species there is an abundance of closely related host species, which enables the parasites to readily undergo host switches and thus reach a high frequency. Host tree #5 (Figures 3C and S2B) shows the opposite extreme: the host tree consists of several clades that are only distantly related to each other. Parasite spread and survival within those clades is difficult because these clades are small, and switches between clades are unlikely. Combined, this explains the low infection frequencies observed on this tree. Host tree #25 (Figure S2C, D) contains a large clade of closely related host species in which the parasites can thrive. If the parasites are successful in infecting this large clade, they can reach a high frequency of infected host species (Figure S2C). However, this clade is very isolated from the other clades and connected to the rest of the tree by a long branch. Therefore, in many cases the parasites fail to reach this clade and are confined to the other, much smaller clades (Figure S2D). As a consequence, we observe a bimodal distribution of infection frequencies for this tree.

To formalise some of the above intuitive explanations for variation in parasite abundance across host trees, we calculated for each host tree the Shannon index for the distribution of host species among different host clades (see SI section 1.3). This Shannon index is greater the more host clades there are and the more evenly species are distributed among those clades. Figure 4 shows that the Shannon index is negatively correlated with the fraction of infected host species, indicating that host trees whose species are clustered in few large clades are most conducive to parasite spread. In line with these results, we also found that tree imbalance, as measured by the Colless index (Colless 1982; Heard 1992), has a similar effect but explains less of the variance in infection frequencies than the Shannon index of clade sizes (see SI section 3.1; Figure S3).

**Figure 4.**
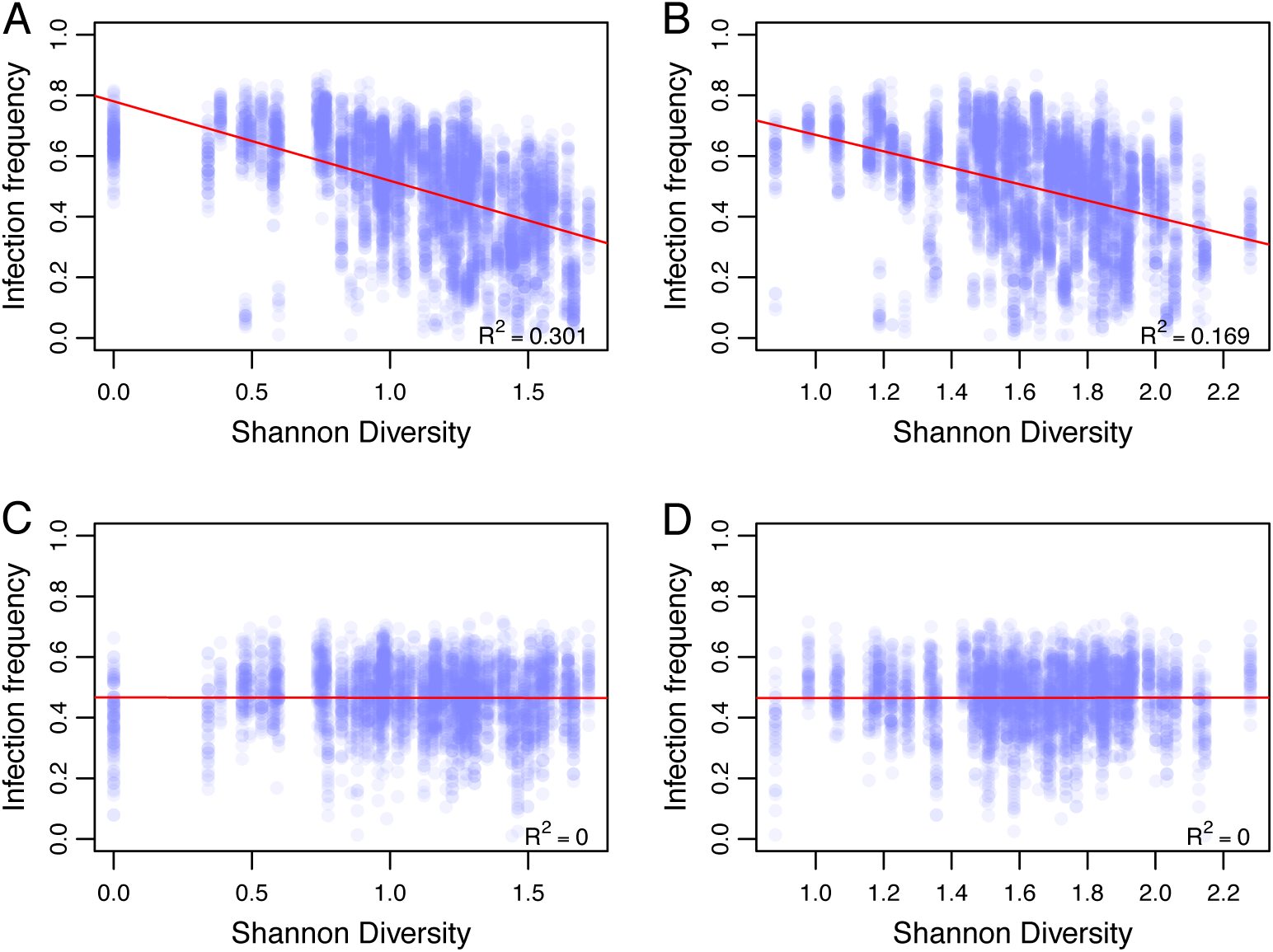
Fraction of infected hosts at the end of simulations against the Shannon index of host species distribution within the respective host tree, with (A,B) or without (C,D) the phylogenetic distance effect. Each dot represents the outcome of a single simulation; simulations in which the parasites became extinct were discarded. Partitioning of host trees into subtrees (or clades) and calculating the Shannon index was performed as described in SI section 1.3, with the height parameter set to either 100 (plots A and C, corresponding to few large subtrees) or 50 (plots B and D, corresponding to more but smaller subtrees). Red lines show the fit of a linear regression with R^2^ values indicated. All parameters take standard values.

### Robustness to parasite parameters and model assumptions

We repeated all simulations with a higher parasite transmission rate (β=1) and a higher extinction rate (*v*=2). Figures S4 and S5 show that our results are very robust to this change in parameters. We also re-ran our simulations relaxing the assumption that no coinfections can occur, that parasites can be lost during host speciation or that they can speciate within a host linage; again, this did not qualitatively affect our results (Figures S6 to S8).

### Host tree size

We next asked how the equilibrium size of the host trees – determined by the carrying capacity *K* – affects the dynamics of parasite spread. In the absence of the phylogenetic distance effect, increasing host tree size results in both an increasing probability of parasite survival and an increasing number of infected hosts at the end of simulations where parasites do survive (Figure 5). Both of these results are straightforward in the light of standard epidemiological models with density-dependent transmission in well-mixed host populations (Keeling & Rohani 2008). In the presence of the phylogenetic distance effect, there is a comparatively modest increase in the parasite survival probability with increasing host tree size, and no change in the infection frequency. This is because from any given infected host species, the number of uninfected hosts that can be reached through host-shifts will generally be limited by the phylogenetic distance effect rather than the total size of the tree.

**Figure 5.**
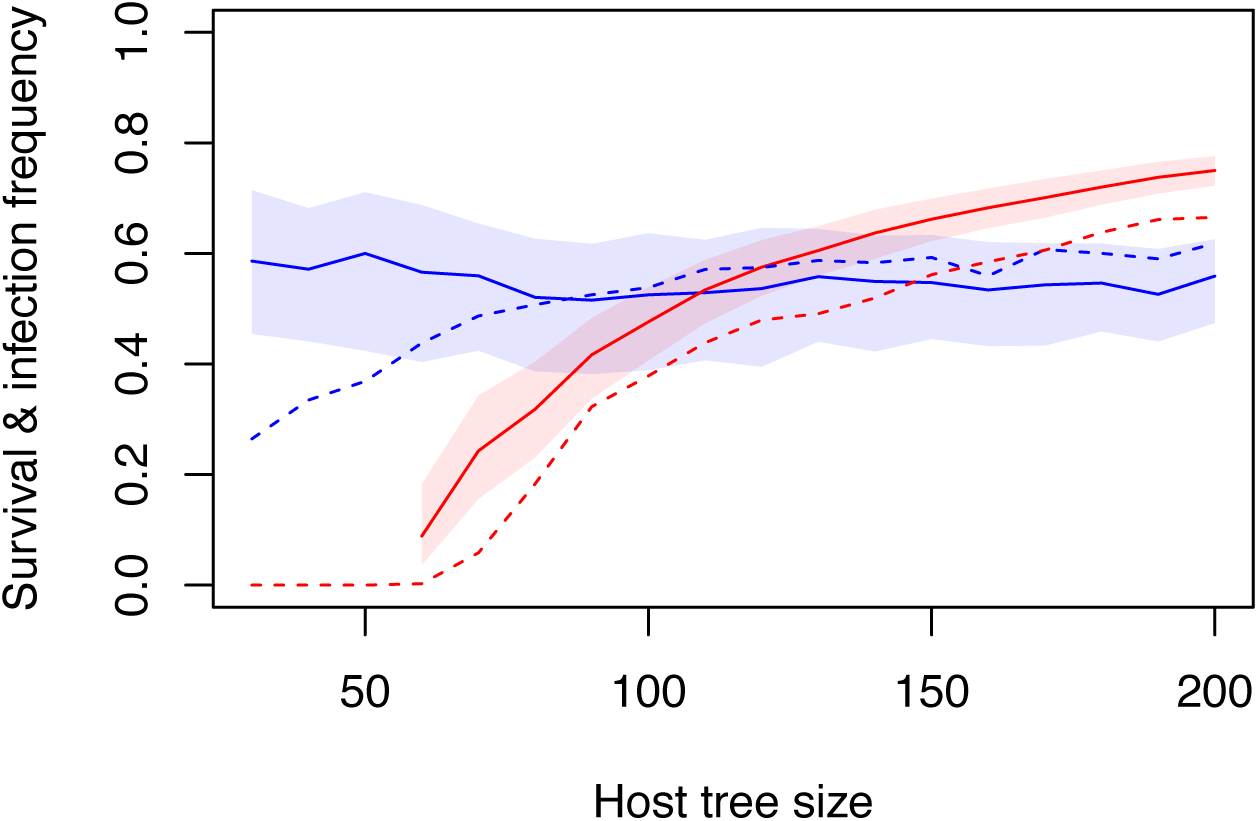
Influence of the equilibrium host tree size on parasite survival rates and infection frequencies in presence (blue) and absence (red) of the phylogenetic distance effect. Dashed lines show the fraction of simulations in which the parasites invaded the host tree and survived until the end of the simulations. Solid lines show the median fraction of infected host species at the end of the simulations for those simulations in which the parasites survived, with shadings indicating the interquartile range. Equilibrium host tree size was modified by varying the carrying capacity parameter *K* over a range of values from 60 to 400. All other parameters take standard values.

### Dynamics of host diversification

The results presented above all assumed that host trees evolved under the same birth-death process, with a speciation rate of *λ*=1 and an extinction rate of μ=0.5. In order to explore the impact of host diversification on parasite spread, we generated sets of host trees with increasing values of *λ* and μ while keeping the difference λ–μ constant. This means that for all sets of host trees generated, the host trees will initially grow at the same net diversification rate but when they reach their carrying capacity, the rate at which new host species are born and go extinct increases (both occurring at rate μ).

Figure 6A shows that in the presence of the phylogenetic distance effect, the host tree sets generated in this way vary strongly in both the parasite survival probability and the fraction of infected host species. When host trees evolve with very low speciation and extinction rates, the parasites almost always become extinct, and if they survive they reach only a very low infection frequency. This is because branches are very long in such host trees, resulting in large phylogenetic distances between host species that are difficult to overcome by the parasites. When λ and μ are high, there will be much turnover in host species and genetic distances will become short so that parasite spread is facilitated, resulting in a high fraction of simulations where parasites survive and reach high infection frequencies.

**Figure 6.**
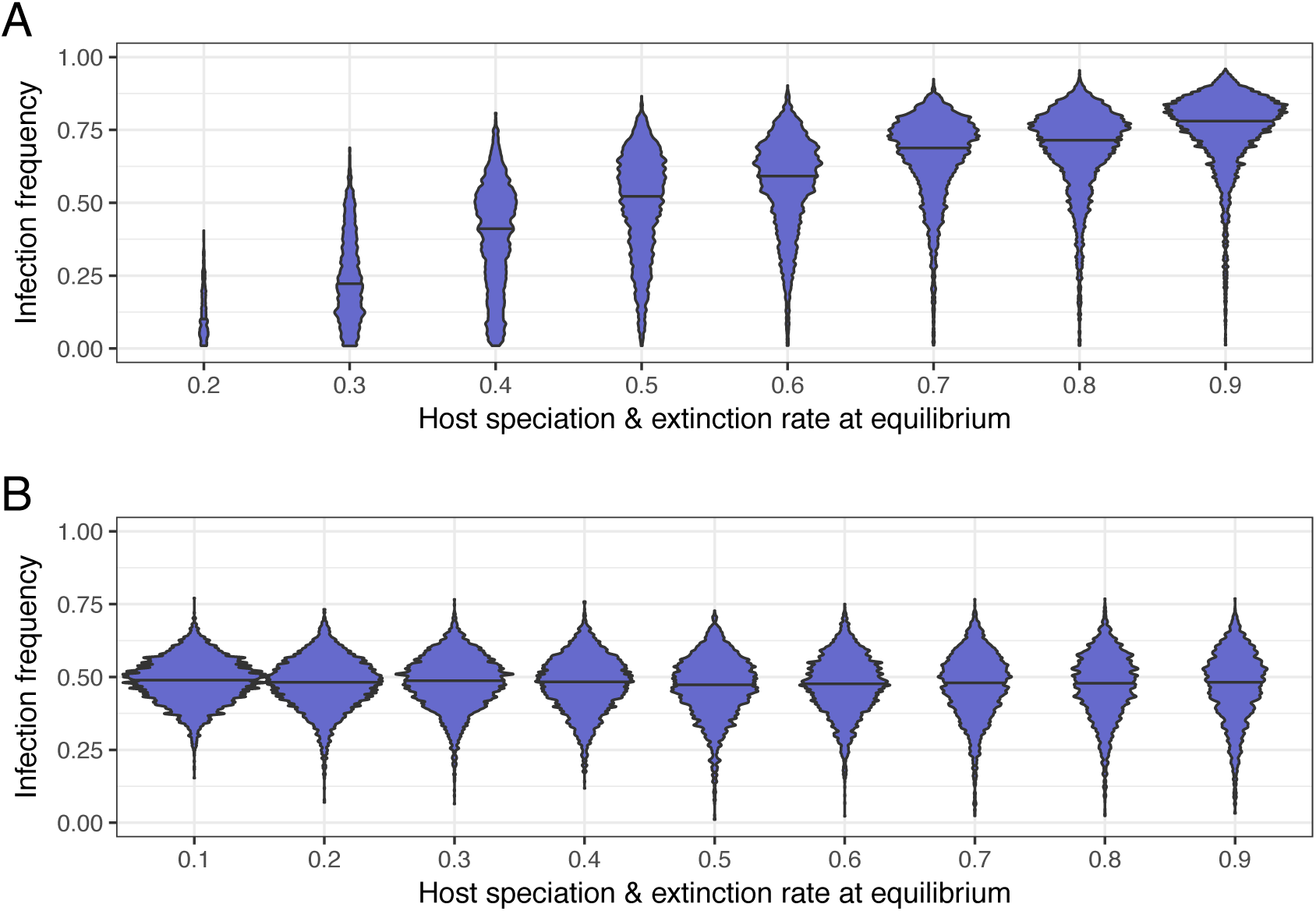
The impact of host speciation and extinction rate at equilibrium on the fraction of infected host species with (A) and without (B) the phylogenetic distance effect. Violins show the distribution of infection frequencies, with the total area of each violin being proportional to the number of simulations where the parasites survived. Equilibrium speciation and extinction rates where varied by using host extinction rates μ ranging from 0.1 to 0.9. At the same time, we varied the host speciation rate *λ* from 0.6 to 1.4 in order to maintain a constant net diversification rate of *λ*–μ=0.5 during the early stages of host evolution. Parasite parameters take standard PDE and no-PDE values.

In the absence of the phylogenetic distance effect, mean infection frequencies are not affected by *λ* and μ (Figure 6B). However, the probability of parasite survival decreases slightly with increasing *λ* and μ. This is because host species numbers vary more through time with high than with low host speciation and extinction rates (results not shown), producing correspondingly strong stochastic variation in infection rates. As a result, when *λ* and μ are high, stochastic parasite extinction is more likely than when *λ* and μ are low.

Finally, we explored whether host net diversification rate (*λ*–μ) or species turnover (μ/*λ*) had any impact on the dynamics of parasite spread beyond the impact of the rate of speciation and extinction in the steady state discussed above. We generated eight additional sets of host trees with different combinations of values for λ–μ and μ/λ (see SI section 1.4). Under the phylogenetic distance effect, the parasite survival rate and the fraction of infected hosts increases with both net diversification rate and host species turnover on these trees (Figure S9A). However, the results are always very similar with identical host extinction rates, suggesting that early host tree evolution was not important. In the absence of the phylogenetic distance effect, different host tree sets only differ mainly in the fraction of simulations where the parasites survived (Figure S9B), presumably again due to different degrees of stochastic fluctuations in host tree size.

## Discussion

Using a mathematical model, we have investigated how the phylogenetic distance effect (preferential host-shifts between closely related species) impacts the prevalence and distribution of parasites across host species. Our model makes a number of predictions: all else being equal and in the presence of the phylogenetic distance effect, 1) host trees in which most species are clustered in a few large clades should harbour more parasites than those consisting of many small clades, 2) host trees characterised by high species turnover (including rapid adaptive radiations) should harbour more parasites than host trees that are evolutionarily more inert, and 3) small and isolated clades within trees should harbour fewer parasites than large clades. These predictions can be tested without any cophylogenetic analyses and indeed, without any knowledge about phylogenetic relationships between the parasites. In contrast to previous models where parasites only switch between extant host species (Engelstädter & Hurst 2006; de Vienne *et al.* 2007; Cuthill & Charleston 2013; Waxman *et al.* 2014), in our model parasite and host diversification occurs concurrently and potentially on similar time scales.

The power of our predictions depends on how strong the phylogenetic distance effect is, both in absolute terms and relative to other effects. The phylogenetic distance effect emerges from the fact that related species tend to be physiologically and immunologically similar, thus increasing the chances that a parasite can successfully replicate in a new host. However, relevant host traits such as the presence or absence of certain cell surface receptors may also evolve repeatedly during host diversification. This can give rise to ‘clade effects’ in which a host clade that is only distantly related to a donor host may nevertheless have a high propensity to be recipients of a parasite (Longdon *et al.* 2011; Waxman *et al.* 2014). Moreover, the probability of host-shifts will depend not only on similarity between host species, but also on opportunities for parasites from one species to encounter hosts from another species. This means that both geographical range overlap and ecological interactions between donor and potential recipient host species may be important determinants of host-shifts. Finally, de Vienne *et al*. (2009) showed that the phylogenetic distance between a native and a new parasite can also be a good predictor of infection success. All of these factors may obscure the phylogenetic distance effect.

Little is known about the relative importance of (phylo)genetic vs. ecological factors for host-shifts, but it appears that this varies widely across systems. On the one hand, several pathogens (e.g., influenza viruses and *Mycobacterium tuberculosis*) have shifted between humans and domesticated animals such as cattle or fowl – species that are only distantly related to humans but have close physical contact (Smith *et al.* 2009; Ren *et al.* 2016). On the other hand, several studies have reported evidence for a strong phylogenetic distance effect. For example, in microalgae-virus associations in the open sea where no ecological barriers to host-shifts should exist, there seems to be a clear signal for the phylogenetic distance effect (Bellec *et al.* 2014). In a study of rabies in bats, host genetic distance was identified as a key factor for host-shifts whereas ecological factors (range overlap and similarities in roost structures) had no predictive power (Faria *et al.* 2013).

The case of *Wolbachia,* an intracellular bacterium infecting nematodes and arthropods (Werren *et al.* 2008), indicates that even for a single parasite there may be considerable variation in the relative importance of different factors affecting host-shift rates. For example, *Wolbachia* underwent preferential host-shifts to related species within the spider genus *Agelenopsis* (Baldo et al. 2008). By contrast, in mushroom-associated dipterans, ecological similarity (mycophagous vs. non-mycophagous) appeared to be an important determinant of *Wolbachia* host-shifts whereas host phylogeny and sympatry did not appear to play a major role (Stahlhut *et al.* 2010). In bees, neither phylogenetic relatedness between hosts nor ecological interactions (kleptoparasitism) predicted *Wolbachia* host-shifts (Gerth *et al.* 2013). Among different orders of arthropods, our prediction that larger clades should have higher infection levels than smaller clades is not supported in *Wolbachia* (Weinert *et al.* 2015), perhaps indicating that at least at this level the phylogenetic distance effect is not important. Overall, the *Wolbachia-*arthropod system is characterised by complex patterns of codiversification that differ between *Wolbachia* strains and host taxa and that we are only beginning to understand (e.g., Gerth *et al.* 2014; Bailly-Bechet *et al.* 2017).

In order to keep our model as simple as possible we made several assumptions. Most importantly, we assumed that each parasite species is strictly associated with a single host species only. This assumption will be met in parasites that are highly specialised on their hosts or that are vertically transmitted, so that transmission between host individuals belonging to different species is very limited. For parasites infecting multiple hosts, we expect that the phylogenetic distance effect should be less pronounced and our results therefore less applicable. Host-shifts were modelled as density-dependent transmission events, i.e. the more host species there are within the host phylogeny, the greater the rate of host-shifts for a parasite. Given that tree size was roughly constant and not affected by the parasites in our model, we again believe that the assumption of density-dependent (as opposed to frequency-dependent) transmission is not crucial to our results. Finally, we assumed an exponential decline in host-shift rates with increasing phylogenetic distance between hosts. This is arguably the simplest function one can assume for this relationship. A sigmoidal relationship has also been proposed (Engelstädter & Hurst 2006) and in a study of RNA viruses in mammals was found to explain the data better than the exponential function (Cuthill & Charleston 2013), but it remains to be seen how general this result is.

In conclusion, we have developed a model of host-parasite codiversification that should be most suitable for parasites that are host-specific and undergo preferential host-shifts according to the phylogenetic distance effect. Our model provides a novel framework to understand host-shift dynamics across large numbers of host species and over long evolutionary time periods. This framework has enabled the generation of several testable predictions regarding the distribution and frequency of parasites, highlighting the importance of host phylogeny in shaping the process of codiversification.

## Acknowledgments

We thank Sylvain Charlat, Ben Longdon, Daniel Ortiz-Barrientos and Tanja Stadler for helpful discussions and Sylvain Charlat, Ben Longdon, Nathan Medd, Damien de Vienne and Lucy Weinert for insightful comments on our manuscript. NF acknowledges funding from an Australian Postgraduate Award and a Global Change Scholars Award from The University of Queensland.

